# Tsetse RNA Virome: Novel Iflavirus Genomes in *Glossina morsitans* and Other Tsetse Species

**DOI:** 10.1101/2021.10.23.465572

**Authors:** M Manni, EM Zdobnov

## Abstract

Tsetse flies (*Glossina* spp.) are vectors of Human and Animal African trypanosomiasis. The tsetse microbiome has been extensively studied in the context of bacterial endosymbiont-host interactions, however, remarkably little is known about the tsetse virome with only one well-characterized DNA virus, the salivary gland hypertrophy virus (SGHV). Here we report the genomes of four iflaviruses identified in tsetse flies and their distribution in public RNA-seq libraries, mainly from laboratory colonies. Strikingly, the iflavirus identified in *G. morsitans*, provisionally named Glossina iflavirus 1 (GliflaV1), is present in all 136 RNA-seq libraries of *G. morsitans* maintained at different institutions, and displays a broad tissue tropism and high abundance, reaching up to 15% of library content. Its ubiquitous distribution and presence in the reproductive tissues, intrauterine larvae, and teneral flies suggest it is part of the initial core microbiota maternally transmitted to the progeny. None of the *G. morsitans* samples harbor iflaviruses identified in the other three, more closely related, tsetse species which, conversely, do not harbor the iflavirus from *G. morsitans*. Though apparently asymptomatic, these infections may influence tsetse host fitness, developmental or biological processes which might be relevant in the context of tsetse population control strategies, mass rearing, and paratransgenesis, and open up new opportunities to study the quadripartite system of interactions among the invertebrate host, the parasitic protozoan, and both viral and bacterial symbionts.

## Introduction

Tsetse flies (Diptera: Glossinidae) can transmit parasitic African trypanosomes causing sleeping sickness in humans and nagana in cattle (Cox 2004). Despite the ongoing efforts to control these diseases, trypanosomiasis still represents a threat in African areas inhabited by tsetse flies (Cecchi et al. 2015; Büscher et al. 2018). The lack of vaccines or medicines to directly control the disease makes the application of vector control strategies an effective measure to reduce disease transmission (Courtin et al. 2015; Lehane et al. 2016). Among these approaches, the sterile insect technique (SIT) and paratransgenesis are promising methods to control tsetse flies, which are viviparous (i.e. the mother supports the development of her progeny in an intrauterine environment) and feed exclusively on vertebrate blood. The additional nutritional requirements essential for biological function are fulfilled by maternally transmitted symbionts which also regulate the host immune system (Wang et al. 2009; Weiss et al. 2011; Snyder et al. 2012; Michalkova et al. 2014; Snyder and Rio 2015; Rio et al.). These bacteria, namely the obligate *Wigglesworthia glossinidia*, the commensal *Sodalis glossinidius* and *Wolbachia* have been extensively studied in the context of symbiont-host interactions and for paratransgenesis applications (Aksoy et al. 2008; De Vooght et al. 2018; Demirbas-Uzel, De Vooght, et al. 2018; Yang et al. 2021). In adult flies, *Wigglesworthia* localization is restricted to the bacteriome organ, where it resides intracellularly within specialized cells (bacteriocytes), and in the female milk glands. This symbiosis is essential for tsetse survival as flies without *Wigglesworthia* are unable to reproduce (Pais et al. 2008), and aposymbiotic flies (i.e. flies that have developed from larvae reared in the absence of the bacterial symbionts) are severely immuno-compromised (Weiss et al. 2012; Wang et al. 2013). *Sodalis* (Snyder et al. 2010) resides extracellularly in various tissues (Balmand et al. 2013) or intracellularly within gut epithelial cells (Wang et al. 2013), and is not required for tsetse survival with its distribution varying in tsetse populations (Kame-Ngasse et al. 2018; Tsagmo Ngoune et al. 2019). Similarly, the distribution of *Wolbachia*, which is mainly found in the reproductive tissues (Balmand et al. 2013; Schneider et al. 2018) and confers strong cytoplasmic incompatibility (CI) during embryogenesis, (Alam et al. 2011) is patchy (Doudoumis et al. 2012).

Though adult tsetse can harbor a more complex microbiota with additional bacterial species, their distribution is highly variable (Geiger et al. 2009; Lindh and Lehane 2011; Aksoy et al. 2014), they are transitory and likely acquired through the environment, as they are absent in teneral flies (newly emerged adults which have not yet fed). Recently, *Spiroplasma* has been described in individuals from various populations (Doudoumis et al. 2017; Schneider et al. 2019), with its frequency and prevalence varying among different *Glossina* species, populations and individuals (Alam et al. 2012; Symula et al. 2013; Aksoy et al. 2014). For example, *Spiroplasma* appears to be absent in species from the *Morsitans* and *Fusca* subgenera, whereas it is found in species of the *Palpalis* group (Doudoumis et al. 2017). Despite the in-depth knowledge of tsetse bacterial endosymbionts, remarkably little is known about the tsetse virome. Only one dsDNA virus, the salivary gland hypertrophy virus (SGHV), has been well-characterized in tsetse flies (Abd-Alla et al. 2008; Demirbas-Uzel, Parker, et al. 2018; Meki et al. 2018; Demirbas-Uzel et al. 2021).

Here we characterized four iflavirus genomes (order *Picornavirales*) and their distribution in *G. morsitans* and other tsetse species (*G. palpalis palpalis, G. palpalis gambiensis, G. pallidipes, G. brevipalpis, G. fuscipes*, and *G. austeni*). Iflaviruses are small viruses with positive-sense single-stranded RNA genomes that typically contain a single open reading frame (ORF) encoding a large polyprotein flanked by 5’ and 3’ untranslated regions (UTRs). Different insect-specific viruses (ISVs) of the *Iflaviridae* family have been described in various arthropod species. They only replicate in insect cells and are maintained in the populations through vertical or horizontal transmission. ISVs can have physiological impacts on their hosts, for example, altering mosquitoes’ susceptibility to arbovirus infection (Nasar et al. 2015; Schultz et al. 2018; Baidaliuk et al.). The tsetse virome might as well impact host fitness, biological and physiological processes, and its characterization may be relevant in the context of paratransgenesis and population control strategies. Characterization of RNA viruses that share a coevolutionary history with tsetse flies opens up exciting windows of opportunities to study the quadripartite interactions occurring among the invertebrate host, the virus, the bacterial endosymbionts, and the protozoan pathogens in this already well-characterized system.

## Materials and methods

### Iflavirus discovery

We collected all 143 publicly available RNA-Seq datasets of *G. morsitans* deposited in the NCBI Sequence Read Archive (SRA) database (Sayers et al. 2020) and converted them to FASTQ files using the NCBI SRA toolkit (https://github.com/ncbi/sra-tools). These libraries represent different tissues from single individuals or pools of flies from different laboratory colonies established from various geographic locations (Supplementary Table 1). Specifically, *G. morsitans* libraries were derived from laboratory colonies of: *i)* Yale (USA), established with puparia from wild samples from Zimbabwe from 1993, representing samples from testes and accessory glands (PRJNA295435) (Scolari et al. 2016), wild-type and aposymbiotic (intrauterine) larvae (PRJNA309164) (Benoit et al. 2017), trypanosome-infected whole midguts (PRJNA314786) (Aksoy et al. 2016), female bacteriome (PRJNA335358) (Bing et al. 2017), antennae and heads (PRJNA343269; PRJNA343267) (Kabaka et al. 2020), infected and uninfected proboscis (PRJNA354110) (Awuoche et al. 2018); infected and uninfected cardia (PRJNA358388) (Vigneron et al. 2018), Whole midgut of aposymbiotic adult flies (PRJNA368970) (Bing et al. 2017); infected and uninfected midguts (PRJNA368987), male reproductive tissues (accessory glands and testes) of aposymbiotic and wild type flies (PRJNA394896), adult male antennae (PRJNA429025), milk from tsetse fly larvae gut (PRJNA429038), carcasses (PRJNA205861) (Benoit et al. 2014), infected and uninfected Salivary glands (PRJNA217801) (Telleria et al. 2014); *ii)* the colony at the Institute of Tropical Medicine Antwerp (ITMA) (Belgium), originated from pupae collected in Kariba (Zimbabwe) and Handeni (Tanzania) (P et al. 1993), consisting of libraries from salivary glands of *T. brucei* -infected and uninfected flies (PRJNA327366) (Matetovici et al. 2016), and abdomens of flies with and without *Sodalis* (PRJNA476840) (Trappeniers et al. 2019); *iii)* the colony maintained at the Tsetse Research and Mass Rearing Facility, Institute of Zoology, Slovak Academy of Sciences, Bratislava, Slovakia, including libraries of ovaries and uterus from virgin and pregnant flies (PRJNA448829) (Procházka et al. 2018); iv) and from pupae maintained in the Department of Biology insectary at West Virginia University that were supplied by the Institute of Zoology, Slovak Academy of Sciences (Bratislava, Slovakia), these libraries includes male and female bacteriomes and midguts (PRJNA668823) (Medina Munoz et al. 2021). Seven libraries assigned to different accession numbers and under different BioProjects appear to be duplicated (i.e. raw libraries have the exact same content). Thus we included only the version associated with more metadata, bringing the total number of libraries down to 136 libraries. The libraries were analyzed with FastQC v0.11.9 (https://www.bioinformatics.babraham.ac.uk/projects/fastqc/) to identify adapters. bbduk from the BBTools suite v38.86 (Bushnell) was used to trim sequencing adapters and low-quality regions, and to filter reads with an average quality below 20. To reduce the computational demand, reads matching to the host genome (*G. morsitans*, assembly accession: GCA_001077435.1) were computationally subtracted from the clean libraries with bbduk using a *k-*mer size of 30, i.e., reads with 31-mers matching the reference were removed. This also excludes from our analyses potential viruses with high sequence similarity to possible endogenous viral elements in the tsetse nuclear genome. The remaining reads were de novo assembled using SPAdes v3.14.1 (Prjibelski et al. 2020) for both the paired-end and single-end libraries using the “--rna” mode with *k*-mers of 21, 33, 55, 77 and 97 for reads >=100 bp or with default parameters for libraries of <100 bp. After excluding short sequences (< 500 bp) and low-entropy sequences using bbduk, the remaining contigs were annotated with the library name from which they were obtained, combined in a single FASTA file and clustered using CD-HIT 4.8.1 (Fu et al. 2012). To identify novel viruses, representative sequences from each cluster were queried against a local BLAST database of virus proteins deposited in the NCBI database. Only clusters with members displaying high sequence coverage and deriving from the majority of libraries and studies (defined as having different SRP accessions) were considered in this analysis.

To verify the quality of assembled sequences, reads from clean libraries in which a putative virus genome was identified were mapped to the corresponding sequence using bbmap v38.86 (Bushnell) (“minid” = 0.96, “maxindel” = 4, “minaveragequality” = 20). Coverage and other mapping statistics were calculated using bbmap. The resulting SAM files were converted to Binary Alignment (BAM) files with Samtools v1.13 (Danecek et al. 2021) and mapping reads were visually inspected using the Integrative Genomics Viewer (IGV) (Thorvaldsdóttir et al. 2013). The mapping results were used to identify variations within the libraries compared to the representative genome using Freebayes v1.3.5 (Garrison and Marth 2012). The Python package DNA Features Viewer (Zulkower and Rosser 2020) was used to visualize annotations and coverage data.

MAFFT (Katoh et al. 2019) was used to compute alignments of coding-complete assemblies obtained from different libraries. To carry out additional analyses, such as gene annotation and phylogenetic analysis, one representative was chosen for highly similar assemblies (>98% nucleotide identity, i.e. considered to belong to the same viral species).

To assess the distribution and abundance of the virus across all samples, adapter and quality filtered libraries were mapped to the representative genome using bbmap as described above. Mapping statistics such as breadth and depth of coverage were calculated using bbmap. The numbers of mapping reads were used to compute RPM (read per million reads) and the percentage of total number of the adapter and quality filtered libraries that were used for mapping. Supplementary Table 1 lists all the SRA run accession numbers along with metadata and mapping statistics.

### Virus genome characterization

Open reading frames (ORFs) were predicted using the NCBI Open Reading Frame Finder (https://www.ncbi.nlm.nih.gov/orffinder/) with a minimum ORF length of 75 nucleotides using the standard genetic code. Predicted protein sequences were compared to the NCBI non-redundant protein database (nr) using BLASTP (Camacho et al. 2009) and analyzed for protein domains using the NCBI Conserved Domain Search Service (Lu et al. 2020), HMMER (https://toolkit.tuebingen.mpg.de/tools/hmmer) and HHPred tool (https://toolkit.tuebingen.mpg.de/#/tools/hhpred). Sequence motifs were searched using the MEME Suite v5.1.1 (Bailey et al. 2015).

### Iflaviruses characterization in other Glossina species

To investigate whether the virus identified in *G. morsitans* or divergent relatives were also present in other *Glossina* species we analyzed the 113 RNA-seq libraries available from other *Glossina* species consisting of samples from laboratory colonies and a smaller number of libraries from wild-caught individuals (Supplementary Table 2). Some of these were duplicated libraries, which we excluded from the analysis retaining the ones associated with more metadata, for a total number of 91 libraries deriving from *G. palpalis gambiensis* (n=26), *G. pallidipes* (n=20), *G. brevipalpis* (n=17), *G. fuscipes* (n=16), *G. palpalis palpalis* (n=10), and *G. austeni* (n=2). Initially, we mapped all libraries to the virus identified in *G. morsitans* using bbmap with loose mapping parameters. Selected libraries with mapping reads were de novo assembled following the strategy as described above. Sequences with similarity to the virus found in *G. morsitans* were retrieved with BLASTn and tBLASTn searches. Candidate assemblies of related but divergent viruses were validated by re-mapping RNA-Seq data to the assemblies to check for sufficient coverage and identify possible misassembly as described above. Representative virus genomes were selected on the basis of sequence coverage and preferentially of libraries derived from individual flies where available. Intra-library polymorphisms were estimated using Freebayes (Garrison and Marth 2012) as described above. Validated assemblies of iflaviruses from different species were compared using reciprocal BLASTn analysis and aligned with MAFFT. Percent identities at nucleotide and protein levels among the novel iflaviruses were calculated from the alignments using clustalO v1.2.4 (“--percent-id” option) (Sievers and Higgins 2014).

To assess the presence of the newly identified viruses across all libraries from all tsetse species, adapter and quality filtered fastq libraries from all species were mapped against the reference of each newly identified virus using bbmap with stricter parameters (“minid” = 0.96, “maxindel” = 4).

### Phylogenetic analysis

Phylogenetic analysis was performed using the polyprotein sequences of the newly-identified *Glossina* viruses and those of known iflaviruses deposited in GenBank. The sequences were aligned using MAFFT employing the G-INS-i algorithm and trimmed with trimAL v3.4.1 (Capella-Gutiérrez et al. 2009) with the option “-automated1”. The best-fit amino acid substitution model was estimated with ModelFinder implemented in IQ-TREE2 (Nguyen et al. 2015), which was used to compute a maximum likelihood phylogenetic tree using 100,000 Ultrafast bootstraps. The phylogenetic tree was visualized using Dendroscope v3.7.6 (Huson and Scornavacca 2012).

### Virus insertions

Endogenous viral elements (EVEs) are common in insect nuclear genomes and can be transcriptionally active. As in our analysis we subtracted reads matching to the corresponding *Glossina* genomes, we do not expect to find nuclear viral insertions corresponding to the newly identified viruses. To further exclude the possibility that the virus contigs are derived from EVEs and to investigate the presence of EVEs from related viruses, all identified iflaviruses and their polyproteins were screened in all the six *Glossina* genomes (*G. morsitans*, GCA_001077435.1; *G. austeni*, GCA_000688735.1; *G. brevipalpis* GCA_000671755.1; *G. fuscipes*, GCA_000671735.1; *G. pallidipes*, GCA_000688715.1; and *G. palpalis*, GCA_000818775.1), using BLASTn and tBLASTn (Camacho et al. 2009).

## Results

### Iflavirus discovery in G. morsitans

We initially identified and characterized the genomic sequence of a positive-sense single-stranded RNA virus in *G. morsitans*. The 136 *G. morsitans* RNA-Seq libraries downloaded from the Sequence Read Archive (SRA) database consist of a total of 5.76 billion reads (∼502 billion bases) and represent different tissues of individual flies or pools from different laboratory colonies (Supplementary Table 1 and Material and Methods section). After removing low-quality reads and reads matching *G. morsitans* genome (assembly accession: GCA_001077435.1), the remaining reads were de novo assembled into contigs. After clustering, we obtained a large group of contigs displaying high sequence similarity (>98%) and, on average, high coverage. Strikingly, assembled contigs in this cluster were derived from almost all *G. morsitans* libraries, indicating that the source of these sequences was very common among tsetse samples. Portions of these contigs shared ∼40% amino identity with polyproteins of positive-sense RNA viruses of the *Iflaviridae* family deposited in NCBI databases. The draft virus assemblies independently assembled from different libraries were validated through remapping and manual inspection. As these genomes were highly similar (>99% identity), we selected one representative genome which displayed high coverage and derived from a single fly library, in order to avoid possible inter-host polymorphism in pooled libraries. The representative genome has a size of 10’749 nt (1’288X depth of coverage, assembled from library SRR8284535) with 32% of GC content and a nucleotide distribution of 33% A, 13% C, 19% G, and 35% U. The predicted length of the open reading frame (ORF) is 9’222 nt and encodes for a single polyprotein of 3’074 aa which harbors the typical protein domains of known iflaviruses (Figure 1).

**Figure 1.**
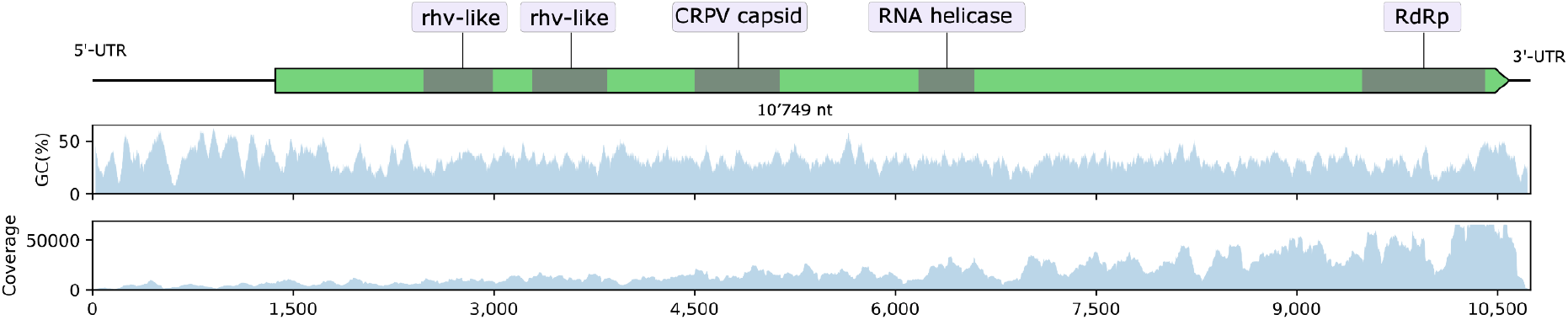
Positive-sense RNA virus identified in *G. morsitans*. (a) Schematic genome organization, GC content, and coverage of Glossina iflavirus 1 (GliflaV1). The virus polyprotein is shown in green and known conserved protein domains in grey. Coverage statistics for all libraries are reported in Supplementary Table 1. Coverage for library SRR2433817 is displayed in the figure.

A portion of the polyprotein shares between ∼30% and ∼40% pairwise amino acid identity to polyproteins of known iflaviruses and 45% identity to the recently discovered Bactrocera tryoni iflavirus 1 virus assembled from a metagenomic analysis of *Bactrocera tryoni* (Diptera: Tephritidae) (Sharpe et al. 2021). Variant analysis on the sample from which the representative genome was derived uncovered 5 SNPs over the ∼10.7 Kbp sequence, which were all synonymous, probably representing a small intra-individual variability. The phylogenetic analysis places the newly discovered virus within the Iflaviridae family with Bactrocera tryoni iflavirus 1 as the known closest virus (Figure 2). On the basis of the phylogenetic evidence and genome organization we tentatively named this virus Glossina iflavirus 1 (GliflaV1).

**Figure 2.**
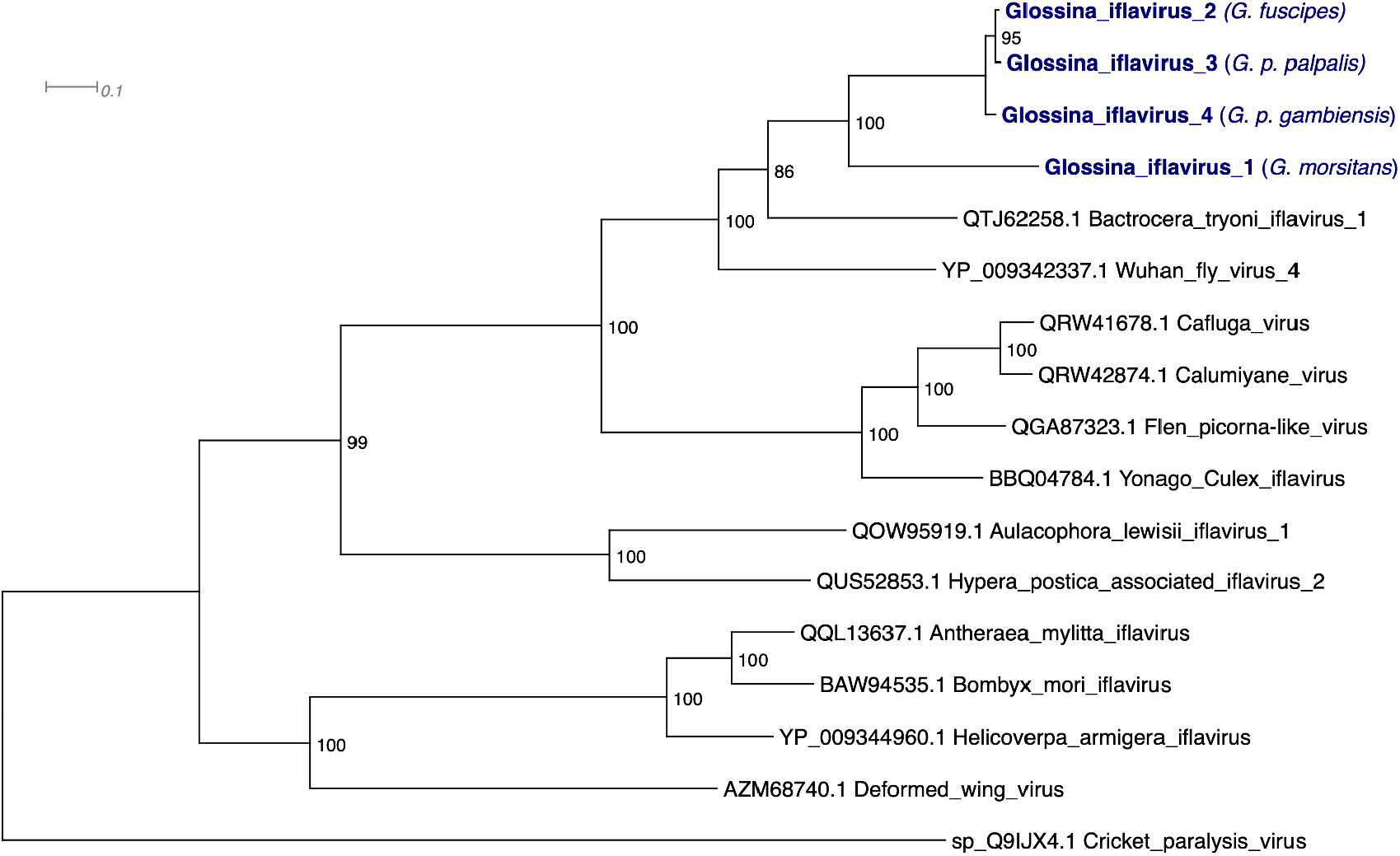
Phylogenetic tree of the newly identified *Glossina* iflaviruses (in blue) with known iflaviruses based on the alignment of the polyprotein sequences. The Cricket paralysis virus (*Cripavirus*) was used as an outgroup. Bootstrap support values are shown on nodes. Branch length indicates the number of amino acid substitutions per site. Host species from which the reference sequence was assembled is reported in parenthesis.

### *Glossina iflavirus 1* is present in all G. morsitans libraries from different laboratory colonies

To define the prevalence and distribution of GliflaV1 in all *G. morsitans* libraries, reads were mapped to the reference genome using strict parameters. Strikingly, GliflaV1 is ubiquitous in all 136 libraries from all independent studies (20/20 BioProject accessions) from different laboratory colonies (and same colony in different periods) maintained at the Yale School of Public health (USA), the Institute of Tropical Medicine Antwerp (Belgium) and the Tsetse Research and Mass Rearing Facility of the Slovak Academy of Sciences (Slovakia) (Supplementary Table 1). GliflaV1 mean depth of coverage varies between 27X (SRR6943612) and 20’285X (SRR2433818) (Supplementary Table 1). The Yale colony was established with puparia of wild samples from Zimbabwe in 1993. The colony at the Institute of Tropical Medicine Antwerp (ITMA) (Belgium), originated from pupae collected in Kariba (Zimbabwe) and Handeni (Tanzania) (P et al. 1993). It is likely that the wild sources of the laboratory strains were infected with GliflaV1 and the virus has been maintained over time likely through vertical transmission.

### Reproductive tract and broad tissue tropism of *Glossina iflavirus 1* in G. morsitans

GliflaV1 is present in samples from a wide range of *G. morsitans* tissues, namely male and female carcasses, bacteriome, midgut, heads, antennae, salivary glands, proboscis, and reproductive tissue from female (ovaries and uterus) and males (testes and male accessory glands) (Figure 3, Supplementary Table 1). The antennal and testes libraries yield the highest depth of coverage (up to approximately 20’000X). Notably, up to 15% of the reads from quality-filtered antennal libraries (including the host reads) is constituted by GliflaV1-derived reads (e.g., SRR4253031: 2’825’863/18’977’315 reads, raw library size: 20’884’277 reads), followed by testis (up to 3.6%) and the bacteriome (up to 3.0%), indicating that this virus can reach abundant levels in these tissues (Figure 3). GliflaV1-derived reads representing more than 1% of the total quality-filtered libraries were observed in other samples/tissues, namely the head, proboscis, carcass, and midgut (Figure 3, Supplementary Table 1). GliflaV1 was also detected in libraries of both wild-type and aposymbiotic intrauterine larvae (PRJNA309164), in milk from the larval gut (PRJNA429038), ovaries, and uterus of virgin and pregnant flies. Given the peculiar reproductive biology of tsetse flies where the larva develops in the mother uterus, the presence of GliflaV1 in the reproductive tracts and developing larvae, suggesting the virus can be vertically transmitted, as it happens for the core bacterial endosymbionts. As the virus appears in all samples, irrespective of the presence of co-infecting microbes (e.g. Sodalis+/-, Trypanosome+/-), and in aposymbiotic flies (flies developed in absence of the resident microbiota, including *Wigglesworthia*) it seems GliflaV1 is not strictly associated with other known tsetse microbes. However, whether the presence of GliflaV1 can influence the composition and abundance of tsetse endosymbionts and pathogens, and/or vice-versa, would warrant further investigation.

**Figure 3.**
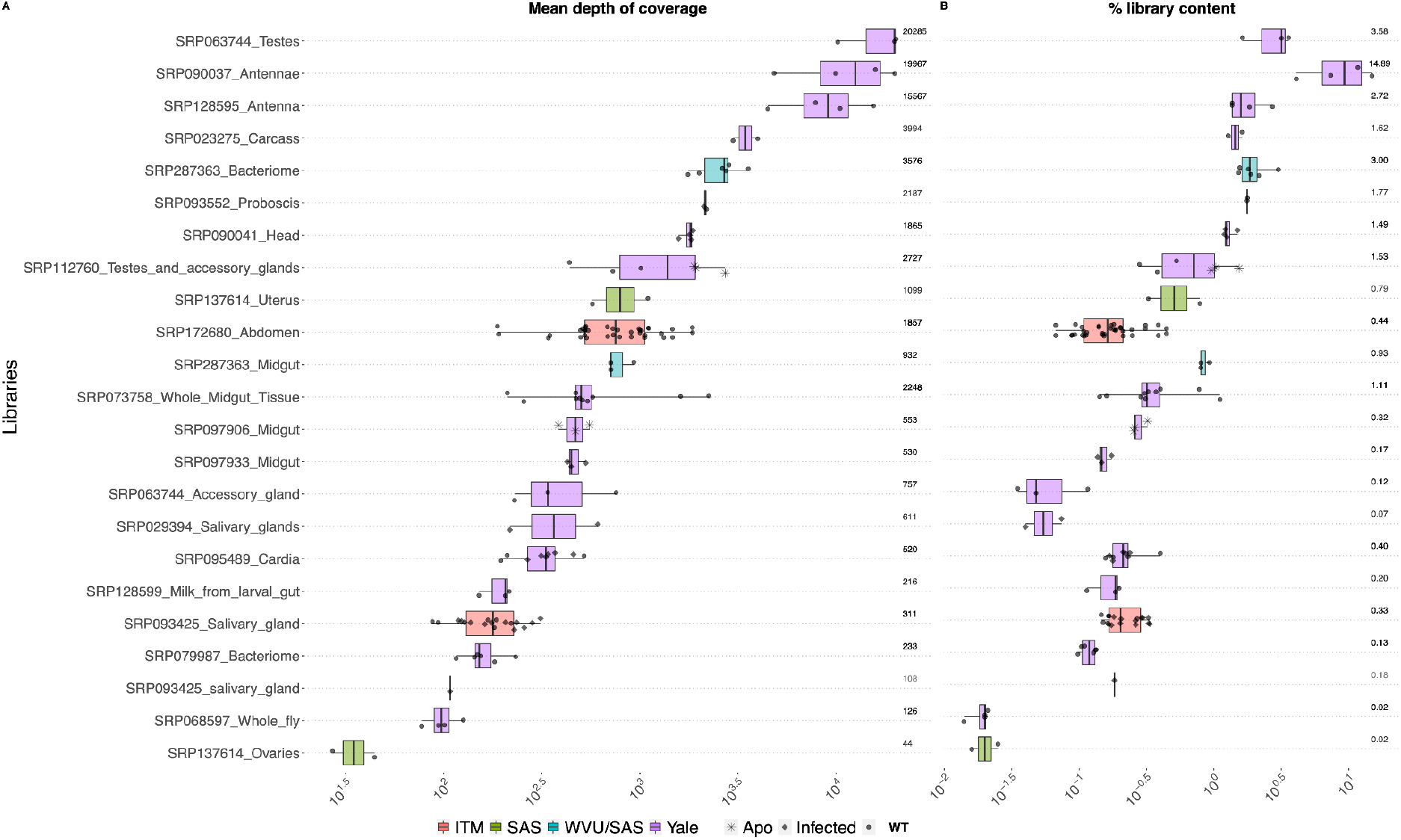
Distribution and abundance of Glossina iflavirus 1 (GliflaV1) in all 136 *G. morsitans* libraries. **a)** Mean depth of coverage to the reference GliflaV1 genome. **b)** Viral abundance as the percentage of virus-derived reads on the total reads from quality-filtered RNA-seq libraries. Libraries are grouped by study (SRP accession) and tissue/developmental stage. For each library group, the max value is displayed on the right of both panels. Colors correspond to different laboratory colonies. Mapping statistics for each library are reported in Supplementary Table 1. Axis are in logarithmic scale. “ITM”, Institute of Tropical Medicine Antwerp (Belgium); “SAS”, Slovak Academy of Sciences, Bratislava, (Slovakia); “WVU/SAS”, West Virginia University with samples supplied by the Slovak Academy of Sciences, Bratislava, (Slovakia); “Yale”, Yale School of Public health (USA). “Apo”, aposymbiotic flies; “Infected”, Trypanosome-infected flies; “WT”, wild-type flies.

### Related but divergent iflaviruses in other Glossina species

To assess whether GliflaV1 could be found in libraries from other tsetse species, we mapped the available 91 RNA-seq libraries from other *Glossina* species (*G. palpalis gambiensis, G. pallidipes, G. brevipalpis, G. fuscipes, G. palpalis palpalis*, and *G. austeni*) to the GliflaV1 genome. None of these libraries contains GliflaV1, except for a few *G. brevipalpis* libraries though at very low abundance (max 2.8X, 72% breadth of coverage). However, diverging reads were mapped on a small fraction of GliflaV1 suggesting that reads from closely related iflaviruses mapped to small conserved regions. We therefore de novo assembled these libraries and obtained coding complete iflavirus genomes from *G. fuscipes, G. p. palpalis* and *G. p. gambiensis*. GliflaV1 shares 59% nucleotide identity with virus from *G. fuscipes*, which we named Glossina iflavirus 2 (GliflaV2, obtained from library SRR650433); 61% nucleotide identity with the one from *G. p. palpalis*, named Glossina iflavirus 3 (GliflaV3, obtained from library SRR5571743); and 60% nucleotide identity with a virus from *G. p. gambiensis*, named Glossina iflavirus 4 (GliflaV4, obtained from library SRR7698158). The similarity among these four *Glossina* iflaviruses ranges approximately from 59% to 88% nucleotide identity, with the iflavirus obtained from *G. morsitans* being the most divergent. On amino acid level, GliflaV1 polyprotein shares 47% identity with the polyprotein of the other three tsetse iflaviruses, with GliflaV3 (from *G. palpalis*) and GliflaV2 (from *G. fuscipes*) sharing the highest protein identity (97%) (Table 1). Importantly, though there are only few libraries from wild-caught flies, GliflaV3 was obtained from a field-collected individual of *G. p. palpalis* (SRR5571743, 930X, 100% coverage), indicating that this group of viruses occur also in natural tsetse populations. Variant analysis on the libraries used for obtaining the representative genomes highlighted no variation for GliflaV3 which was derived from a single fly. Whereas, there were 44 and 43 SNPs for GliflaV2 and GliflaV4, respectively. These libraries are also derived from single adult females, suggesting this variation represents intra-host diversity. For the assembled sequences we adopted a consensus majority rule, i.e. the most frequent allele was kept in the references.

**Table 1.** Pairwise sequence identity at nucleotide (above diagonal) and amino acid (below diagonal) level among the characterized *Glossina* iflaviruses.

|  | GliflaV1 | GliflaV2 | GliflaV3 | GliflaV4 |
| --- | --- | --- | --- | --- |
| GliflaV1 (*G. morsitans*) | - | 58.9% | 61.1% | 60.0% |
| GliflaV2 (*G. fuscipes*) | 51.6% | - | 88.8% | 85.2% |
| GliflaV3 (*G. p. palpalis*) | 52.7% | 97.5% | - | 85.0% |
| GliflaV4 (*G. p gambiensis*) | 51.9% | 92.8% | 93.5% | - |

### Distribution of iflaviruses in Glossina species

To determine the incidence and abundance of all four iflaviruses among all tsetse species, we mapped all libraries against the four iflavirus assemblies (Supplementary Table 2). GliflaV1 identified in *G. morsitans* is absent from species of the *Palpalis* group (*G. palpalis* and *G. fuscipes*; 0/24 libraries) (Table 2). Viceversa, the ones identified from the *Palpalis* group were not found in the *Morsitans* sub-genera (*G. m. morsitans, G. pallidipes, G. austeni*) except for some *G. pallidipes* samples that mapped to GliflaV4, though at low coverage (5.8X; 87% coverage) and with a high proportion of SNPs, suggesting these reads might derive from a closely related but more differentiated virus. GliflaV4 was present in all libraries of *G. p. gambiensis* (26/26 libraries) from Yale University (with samples derived from the Slovak Academy of Sciences colony), and the IRD-CIRAD laboratory colony (PRJNA260242) which were established from flies collected in different areas of Burkina Faso (Hamidou Soumana et al. 2014; Hamidou Soumana et al. 2015). GliflaV4-derived reads account for up to 7% of the quality-filtered library content (approximately 4.8M/67.8M reads of library SRR7698161 from an adult male sample). *G. fuscipes* libraries mapped to GliflaV2 at high abundance in two libraries (max coverage 7’601X, 100% breadth of coverage) from whole bodies of lactating and non-lactating females (SRR7698165 and SRR7698164), whereas few other libraries partially mapped to GliflaV4 at very low coverage. In *G. brevipalpis* libraries we identified a low number of reads partially mapping to GliflaV1 but with low coverage (max 2.8X; breadth of coverage of 73%). No reads mapping to any of the assessed iflaviruses were detected in libraries of *G. austeni*, though only 2 libraries are available. The phylogenetic tree in figure 2 shows the three viruses identified in the Palpalis group (*G. p. palpalis, G. p*.*gambiensis* and *G. fuscipes)* as being more closely related to each other compared to GliflaV1 from *G. morsitans*, suggesting a co-evolutionary history of these viruses within the *Glossina* genus, though this warrants further investigation with large samples from tsetse natural populations and sampling of other *Glossina* species. This virus-host species pattern is also observed in colonies of different species that are co-housed in the same facility where flies might have higher chances to potentially exchange viruses horizontally via the environment.

**Table 2.** Libraries of different tsetse species with reads mapping to the newly identified iflaviruses. Values correspond to the number of virus-positive RNA-seq libraries. The total number of libraries and BioProjects for each species is shown in parenthesis.

|  | GliflaV1 | GliflaV2 | GliflaV3 | GliflaV4 |
| --- | --- | --- | --- | --- |
| *G. morsitans* (n=136; 20 BioProjects) | 136 | - | - | - |
| *G. p. palpalis* (n=10; 1 BioProject) | - | - | 1* | - |
| *G. p gambiensis* (n=26; 3 BioProjects) | - | 11 | 5 | 26 |
| *G. fuscipes* (n=16; 3 BioProjects) | - | 2 | - | - |
| *G. pallidipes* (n=20; 4 BioProjects) | - | - | - | 2(L) |
| *G. brevipalpis* (n=17; 4 BioProjects) | - | - | - | - |
| *G. austeni* (n=2; 1 BioProject) | - | - | - | - |

Note: Libraries with reads mapping to a newly identified iflavivirus but with a breadth of coverage <80% or a depth of coverage <2X are not reported as virus-positive in this table. The full mapping statistics along with metadata such as library characteristics are reported in Supplementary Tables 1 and 2. Libraries with a depth of coverage between 2X and 10X are labelled with (L).
\*field collected

### Genomes of six Glossina species do not harbor insertions related to the novel iflaviruses

Endogenous viral elements have been described in several insect genomes. Because we used the corresponding nuclear genomes for each species to filter host-specific reads prior to the de novo assembly, we do not expect to find insertions of these newly identified iflaviruses, as their reads would have been filtered out before de novo assembly. Nevertheless, tsetse genomes may harbor sequences from ancient insertion events or from related iflaviruses, diverging from the assembled iflaviruses. With homology searches using BLASTn and TBLASTn we did not find regions with homology to these iflaviruses in any of five tsetse genomes indicating they do not harbor insertions from these or closely related iflaviruses.

## Discussion

Bacterial endosymbionts of tsetse flies have been shown to greatly impact host physiology and have been extensively studied. On the contrary, remarkably little is known about the tsetse virome, with only one well-characterized DNA virus, the Salivary Gland Hypertrophy (SGH) virus. We have identified and characterized positive-sense single-stranded RNA viruses that belong to the *Iflaviridae* family in different tsetse species, and assessed their distribution in public available RNA-seq libraries. Insect-specific viruses (ISVs) from this viral family have been described in various insects. In *G. morsitans* we have characterized an iflavirus which we provisionally named Glossina iflavirus 1 (GliflaV1). Its ubiquitous presence in all *G. morsitans* libraries (136/136 libraries from 19 BioProjects) from different colonies strongly indicates that this viral infection is very common and persistent, at least in laboratory colonies. The presence of diverging but related viruses in other *Glossina* species and the overall distribution pattern suggest this virus group shares a common evolutionary history within tsetse species and may have differentiated along with their hosts. Importantly, we found high levels of one virus (GliflaV3) in a field-collected fly of *G. p. palpalis* from the southern region of Cameroon (Tsagmo Ngoune et al. 2017), indicating these infections can occur in wild flies. However, only a limited number of libraries are available from wild-caught individuals and further assessments are required to understand the distribution and prevalence of these viruses in natural populations. This would also allow to untangle the co-evolutionary history of these viruses among *Glossina* species.

It is likely that the source samples used to establish the tsetse colonies were infected with these iflaviruses, which have been maintained across generations likely through vertical transmission, as shown for other ISVs. The vertical transmission is corroborated by the presence of GliflaV1 in the reproductive tracts of *G. morsitans*, including the testes, accessory glands, ovaries, and uterus. In tsetse flies, the larva develops within the uterus where it receives nutrients through the milk glands (Ma and Denlinger 1974). Tsetse *Wigglesworthia* and *Sodalis* endosymbionts are maternally transmitted to the progeny and initially colonize the gut of the developing intrauterine larva (<u>Attardo et al., 2008</u>) through these female milk secretions during the lactation process (Attardo et al. 2008; Balmand et al. 2013). Indeed, the intrauterine larvae and teneral flies harbor a simple and core microbiota that is not yet perturbed by microbes introduced later through feeding and the environment (Medina Munoz et al. 2021). GliflaV1 is present in various tissues (e.g. testis, accessory glands, antennae, gut, and bacteriome) of teneral flies (i.e. flies that have not received the first blood meal) (Supplementary Table 1), and in the milk from the intrauterine larval gut (e.g. SRR6450984), suggesting this virus is part of the initial core microbiome transmitted early to the progeny by vertical transmission. In this phase, flies have the highest susceptibility to trypanosome infections upon acquisition of their first blood meal (Welburn and Maudlin 1992; Haines 2013) also due to the immaturity of the peritrophic matrix (PM) (Lehane and Msangi 1991; Weiss et al. 2014), which forms a protective semipermeable barrier within the intestinal tract. As GliflaV1 appears to be part of the initial core microbiota of the teneral flies, it would be worth investigating whether its presence/absence or abundance might play a role in determining the susceptibility to trypanosomes infections, e.g. by influencing the state and integrity of the PM. Whether the distribution and prevalence of this group of iflaviruses in different *Glossina* species might have an effect on their differential susceptibilities to trypanosomes would also warrant further investigations.

In tsetse, the male ejaculate is assembled into a capsule-like spermatophore structure visible post-copulation in the female uterus (Scolari et al. 2016). Among other components, male seminal fluid proteins which are transferred during copulation can modulate female reproductive physiology and behavior, e.g. impacting sperm storage/use, ovulation, and remating (Scolari et al. 2016; Savini et al. 2021). Given the high abundance of GliflaV1 in testes, with up to 3.6% of the library consisting of GliflaV1-derived reads (e.g. SRR2433817, SRR2424404), it is likely the virus and its protein products can be transferred to females during copulation. Besides the reproductive tracts, GliflaV1 is detected in the bacteriome, which raises the question of whether and how the virus interacts with *Wigglesworthia*, and in libraries from the salivary glands, antennae and heads. Levels of GliflaV1 are particularly abundant (up to 15% of the library content) in the antennal tissue where it might impact the olfaction process and, consequently, the tsetse host-seeking behavior.

The effects on host physiology and biology of these iflaviruses remain to be determined and warrant further investigations as they may potentially impact traits that are relevant for population control strategies, mass rearing, and paratransgenesis. Besides potential practical implications, the characterization of the tsetse RNA virome offers the opportunity to study the quadripartite system of interactions among the viral and bacterial endosymbionts, the arthropod hosts, and the human pathogens.

---

**Supplementary Table 1**. List of all RNA-seq libraries of *G. morsitans* along with mapping statistics, relevant metadata, and corresponding publications.

**Supplementary Table 2**. List of all RNA-seq libraries from the other tsetse species (*G. palpalis gambiensis, G. pallidipes, G. brevipalpis, G. fuscipes, G. palpalis palpalis*, and *G. austeni*) along with mapping statistics, relevant metadata, and corresponding publications.

## Data availability

The genome assemblies of the *Glossina* iflaviruses have been submitted to the NCBI Third Party Annotation database.

## References

Abd-Alla AMM, Cousserans F, Parker AG, Jehle JA, Parker NJ, Vlak JM, Robinson AS, Bergoin M. 2008. Genome analysis of a Glossina pallidipes salivary gland hypertrophy virus reveals a novel, large, double-stranded circular DNA virus. J Virol 82:4595–4611.

Aksoy E, Telleria EL, Echodu R, Wu Y, Okedi LM, Weiss BL, Aksoy S, Caccone A. 2014. Analysis of Multiple Tsetse Fly Populations in Uganda Reveals Limited Diversity and Species-Specific Gut Microbiota. Applied and Environmental Microbiology 80:4301–4312.

Aksoy E, Vigneron A, Bing X, Zhao X, O’Neill M, Wu Y, Bangs JD, Weiss BL, Aksoy S. 2016. Mammalian African trypanosome VSG coat enhances tsetse’s vector competence. PNAS 113:6961–6966.

Aksoy S, Weiss B, Attardo G. 2008. Paratransgenesis Applied for Control of Tsetse Transmitted Sleeping Sickness. In: Aksoy S, editor. Transgenesis and the Management of Vector-Borne Disease. Advances in Experimental Medicine and Biology. New York, NY: Springer. p. 35–48. Available from: https://doi.org/10.1007/978-0-387-78225-6_3

Alam U, Hyseni C, Symula RE, Brelsfoard C, Wu Y, Kruglov O, Wang J, Echodu R, Alioni V, Okedi LM, et al. 2012. Implications of Microfauna-Host Interactions for Trypanosome Transmission Dynamics in Glossina fuscipes fuscipes in Uganda. Applied and Environmental Microbiology 78:4627–4637.

Alam U, Medlock J, Brelsfoard C, Pais R, Lohs C, Balmand S, Carnogursky J, Heddi A, Takac P, Galvani A, et al. 2011. Wolbachia Symbiont Infections Induce Strong Cytoplasmic Incompatibility in the Tsetse Fly Glossina morsitans. PLOS Pathogens 7:e1002415.

Attardo GM, Lohs C, Heddi A, Alam UH, Yildirim S, Aksoy S. 2008. Analysis of milk gland structure and function in Glossina morsitans: Milk protein production, symbiont populations and fecundity. Journal of Insect Physiology 54:1236–1242.

Awuoche EO, Weiss BL, Mireji PO, Vigneron A, Nyambega B, Murilla G, Aksoy S. 2018. Expression profiling of Trypanosoma congolense genes during development in the tsetse fly vector Glossina morsitans morsitans. Parasites & Vectors 11:380.

Baidaliuk A, Miot EF, Lequime S, Moltini-Conclois I, Delaigue F, Dabo S, Dickson LB, Aubry F, Merkling SH, Cao-Lormeau V-M, et al. Cell-Fusing Agent Virus Reduces Arbovirus Dissemination in Aedes aegypti Mosquitoes In Vivo. Journal of Virology 93:e00705–19.

Bailey TL, Johnson J, Grant CE, Noble WS. 2015. The MEME Suite. Nucleic Acids Research 43:W39–W49.

Balmand S, Lohs C, Aksoy S, Heddi A. 2013. Tissue distribution and transmission routes for the tsetse fly endosymbionts. Journal of Invertebrate Pathology 112:S116–S122.

Benoit JB, Attardo GM, Michalkova V, Krause TB, Bohova J, Zhang Q, Baumann AA, Mireji PO, Takáč P, Denlinger DL, et al. 2014. A Novel Highly Divergent Protein Family Identified from a Viviparous Insect by RNA-seq Analysis: A Potential Target for Tsetse Fly-Specific Abortifacients. PLOS Genetics 10:e1003874.

Benoit JB, Vigneron A, Broderick NA, Wu Y, Sun JS, Carlson JR, Aksoy S, Weiss BL. 2017. Symbiont-induced odorant binding proteins mediate insect host hematopoiesis.Lemaître B, editor. eLife 6:e19535.

Bing X, Attardo GM, Vigneron A, Aksoy E, Scolari F, Malacrida A, Weiss BL, Aksoy S. 2017. Unravelling the relationship between the tsetse fly and its obligate symbiont Wigglesworthia: transcriptomic and metabolomic landscapes reveal highly integrated physiological networks. Proceedings of the Royal Society B: Biological Sciences 284:20170360.

Büscher P, Bart J-M, Boelaert M, Bucheton B, Cecchi G, Chitnis N, Courtin D, Figueiredo LM, Franco J-R, Grébaut P, et al. 2018. Do Cryptic Reservoirs Threaten Gambiense-Sleeping Sickness Elimination? Trends in Parasitology 34:197–207.

Bushnell B. BBMap. Available from: sourceforge.net/projects/bbmap/

Camacho C, Coulouris G, Avagyan V, Ma N, Papadopoulos J, Bealer K, Madden TL. 2009. BLAST+: architecture and applications. BMC Bioinformatics 10:421.

Capella-Gutiérrez S, Silla-Martínez JM, Gabaldón T. 2009. trimAl: a tool for automated alignment trimming in large-scale phylogenetic analyses. Bioinformatics 25:1972–1973.

Cecchi G, Paone M, Argilés Herrero R, Vreysen MJB, Mattioli RC. 2015. Developing a continental atlas of the distribution and trypanosomal infection of tsetse flies (Glossina species). Parasites & Vectors 8:284.

Courtin F, Camara M, Rayaisse J-B, Kagbadouno M, Dama E, Camara O, Traoré IS, Rouamba J, Peylhard M, Somda MB, et al. 2015. Reducing Human-Tsetse Contact Significantly Enhances the Efficacy of Sleeping Sickness Active Screening Campaigns: A Promising Result in the Context of Elimination. PLOS Neglected Tropical Diseases 9:e0003727.

Cox FEG. 2004. History of sleeping sickness (African trypanosomiasis). Infectious Disease Clinics of North America 18:231–245.

Danecek P, Bonfield JK, Liddle J, Marshall J, Ohan V, Pollard MO, Whitwham A, Keane T, McCarthy SA, Davies RM, et al. 2021. Twelve years of SAMtools and BCFtools. GigaScience [Internet] 10. Available from: https://doi.org/10.1093/gigascience/giab008

De Vooght L, Van Keer S, Van Den Abbeele J. 2018. Towards improving tsetse fly paratransgenesis: stable colonization of Glossina morsitans morsitans with genetically modified Sodalis. BMC Microbiology 18:165.

Demirbas-Uzel G, Augustinos AA, Doudoumis V, Parker AG, Tsiamis G, Bourtzis K, Abd-Alla AMM. 2021. Interactions Between Tsetse Endosymbionts and Glossina pallidipes Salivary Gland Hypertrophy Virus in Glossina Hosts. Front Microbiol 12:653880.

Demirbas-Uzel G, De Vooght L, Parker AG, Vreysen MJB, Mach RL, Van Den Abbeele J, Abd-Alla AMM. 2018. Combining paratransgenesis with SIT: impact of ionizing radiation on the DNA copy number of Sodalis glossinidius in tsetse flies. BMC Microbiology 18:160.

Demirbas-Uzel G, Parker AG, Vreysen MJB, Mach RL, Bouyer J, Takac P, Abd-Alla AMM. 2018. Impact of Glossina pallidipes salivary gland hypertrophy virus (GpSGHV) on a heterologous tsetse fly host, Glossina fuscipes fuscipes. BMC Microbiology 18:161.

Doudoumis V, Blow F, Saridaki A, Augustinos A, Dyer NA, Goodhead I, Solano P, Rayaisse J-B, Takac P, Mekonnen S, et al. 2017. Challenging the Wigglesworthia, Sodalis, Wolbachia symbiosis dogma in tsetse flies: Spiroplasma is present in both laboratory and natural populations. Sci Rep 7:1–13.

Doudoumis V, Tsiamis G, Wamwiri F, Brelsfoard C, Alam U, Aksoy E, Dalaperas S, Abd-Alla A, Ouma J, Takac P, et al. 2012. Detection and characterization of Wolbachia infections in laboratory and natural populations of different species of tsetse flies (genus Glossina). BMC Microbiology 12:S3.

Fu L, Niu B, Zhu Z, Wu S, Li W. 2012. CD-HIT: accelerated for clustering the next-generation sequencing data. Bioinformatics 28:3150–3152.

Garrison E, Marth G. 2012. Haplotype-based variant detection from short-read sequencing. arXiv:1207.3907 [q-bio] [Internet]. Available from: http://arxiv.org/abs/1207.3907

Geiger A, Fardeau M-L, Grebaut P, Vatunga G, Josénando T, Herder S, Cuny G, Truc P, Ollivier B. 2009. First isolation of Enterobacter, Enterococcus, and Acinetobacter spp. as inhabitants of the tsetse fly (Glossina palpalis palpalis) midgut. Infection, Genetics and Evolution 9:1364–1370.

Haines LR. 2013. Examining the tsetse teneral phenomenon and permissiveness to trypanosome infection. Frontiers in Cellular and Infection Microbiology 3:84.

Hamidou Soumana I, Klopp C, Ravel S, Nabihoudine I, Tchicaya B, Parrinello H, Abate L, Rialle S, Geiger A. 2015. RNA-seq de novo Assembly Reveals Differential Gene Expression in Glossina palpalis gambiensis Infected with Trypanosoma brucei gambiense vs. Non-Infected and Self-Cured Flies. Front Microbiol 6:1259.

Hamidou Soumana I, Loriod B, Ravel S, Tchicaya B, Simo G, Rihet P, Geiger A. 2014. The transcriptional signatures of Sodalis glossinidius in the Glossina palpalis gambiensis flies negative for Trypanosoma brucei gambiense contrast with those of this symbiont in tsetse flies positive for the parasite: Possible involvement of a Sodalis-hosted prophage in fly Trypanosoma refractoriness? Infection, Genetics and Evolution 24:41–56.

Huson DH, Scornavacca C. 2012. Dendroscope 3: An Interactive Tool for Rooted Phylogenetic Trees and Networks. Systematic Biology 61:1061–1067.

Kabaka JM, Wachira BM, Mang’era CM, Rono MK, Hassanali A, Okoth SO, Oduol VO, Macharia RW, Murilla GA, Mireji PO. 2020. Expansions of chemosensory gene orthologs among selected tsetse fly species and their expressions in Glossina morsitans morsitans tsetse fly. PLOS Neglected Tropical Diseases 14:e0008341.

Kame-Ngasse GI, Njiokou F, Melachio-Tanekou TT, Farikou O, Simo G, Geiger A. 2018. Prevalence of symbionts and trypanosome infections in tsetse flies of two villages of the “Faro and Déo” division of the Adamawa region of Cameroon. BMC Microbiology 18:159.

Katoh K, Rozewicki J, Yamada KD. 2019. MAFFT online service: multiple sequence alignment, interactive sequence choice and visualization. Briefings in Bioinformatics 20:1160–1166.

Lehane M, Alfaroukh I, Bucheton B, Camara M, Harris A, Kaba D, Lumbala C, Peka M, Rayaisse J-B, Waiswa C, et al. 2016. Tsetse Control and the Elimination of Gambian Sleeping Sickness. PLOS Neglected Tropical Diseases 10:e0004437.

Lehane MJ, Msangi AR. 1991. Lectin and peritrophic membrane development in the gut of Glossina m.morsitans and a discussion of their role in protecting the fly against trypanosome infection. Medical and Veterinary Entomology 5:495–501.

Lindh JM, Lehane MJ. 2011. The tsetse fly Glossina fuscipes fuscipes (Diptera: Glossina) harbours a surprising diversity of bacteria other than symbionts. Antonie van Leeuwenhoek 99:711–720.

Lu S, Wang J, Chitsaz F, Derbyshire MK, Geer RC, Gonzales NR, Gwadz M, Hurwitz DI, Marchler GH, Song JS, et al. 2020. CDD/SPARCLE: the conserved domain database in 2020. Nucleic Acids Research 48:D265–D268.

Ma W-C, Denlinger DL. 1974. Secretory discharge and microflora of milk gland in tsetse flies. Nature 247:301–303.

Matetovici I, Caljon G, Van Den Abbeele J. 2016. Tsetse fly tolerance to T. brucei infection: transcriptome analysis of trypanosome-associated changes in the tsetse fly salivary gland. BMC Genomics 17:971.

Medina Munoz M, Brenner C, Richmond D, Spencer N, Rio RVM. 2021. The holobiont transcriptome of teneral tsetse fly species of varying vector competence. BMC Genomics 22:400.

Meki IK, Kariithi HM, Parker AG, Vreysen MJB, Ros VID, Vlak JM, van Oers MM, Abd-Alla AMM. 2018. RNA interference-based antiviral immune response against the salivary gland hypertrophy virus in Glossina pallidipes. BMC Microbiology 18:170.

Michalkova V, Benoit JB, Weiss BL, Attardo GM, Aksoy S. 2014. Vitamin B6 Generated by Obligate Symbionts Is Critical for Maintaining Proline Homeostasis and Fecundity in Tsetse Flies. Applied and Environmental Microbiology 80:5844–5853.

Nasar F, Erasmus JH, Haddow AD, Tesh RB, Weaver SC. 2015. Eilat virus induces both homologous and heterologous interference. Virology 484:51–58.

Nguyen L-T, Schmidt HA, von Haeseler A, Minh BQ. 2015. IQ-TREE: A Fast and Effective Stochastic Algorithm for Estimating Maximum-Likelihood Phylogenies. Molecular Biology and Evolution 32:268–274.

P E, Van HJ, De LE. 1993. The history and breeding conditions of three glossine lines (Diptera, Glossinidae) maintained at the Prince Leopold Institute of Tropical Medicine in Antwerp. Revue de Zoologie Africaine [Internet].

Pais R, Lohs C, Wu Y, Wang J, Aksoy S. 2008. The Obligate Mutualist Wigglesworthia glossinidia Influences Reproduction, Digestion, and Immunity Processes of Its Host, the Tsetse Fly. Appl Environ Microbiol 74:5965–5974.

Prjibelski A, Antipov D, Meleshko D, Lapidus A, Korobeynikov A. 2020. Using SPAdes De Novo Assembler. Current Protocols in Bioinformatics 70:e102.

Procházka E, Michalková V, Daubnerová I, Roller L, Klepsatel P, Žitňan D, Tsiamis G, Takáč P. 2018. Gene expression in reproductive organs of tsetse females – initial data in an approach to reduce fecundity. BMC Microbiol 18:144.

Rio RVM, Jozwick AKS, Savage AF, Sabet A, Vigneron A, Wu Y, Aksoy S, Weiss BL. Mutualist-Provisioned Resources Impact Vector Competency. mBio 10:e00018–19.

Savini G, Scolari F, Ometto L, Rota-Stabelli O, Carraretto D, Gomulski LM, Gasperi G, Abd-Alla AMM, Aksoy S, Attardo GM, et al. 2021. Viviparity and habitat restrictions may influence the evolution of male reproductive genes in tsetse fly (Glossina) species. BMC Biology 19:211.

Sayers EW, Beck J, Bolton EE, Bourexis D, Brister JR, Canese K, Comeau DC, Funk K, Kim S, Klimke W, et al. 2020. Database resources of the National Center for Biotechnology Information. Nucleic Acids Res 49:D10–D17.

Schneider DI, Parker AG, Abd-alla AM, Miller WJ. 2018. High-sensitivity detection of cryptic Wolbachia in the African tsetse fly (Glossina spp.). BMC Microbiology 18:140.

Schneider DI, Saarman N, Onyango MG, Hyseni C, Opiro R, Echodu R, O’Neill M, Bloch D, Vigneron A, Johnson TJ, et al. 2019. Spatio-temporal distribution of Spiroplasma infections in the tsetse fly (Glossina fuscipes fuscipes) in northern Uganda. PLOS Neglected Tropical Diseases 13:e0007340.

Schultz MJ, Frydman HM, Connor JH. 2018. Dual Insect specific virus infection limits Arbovirus replication in Aedes mosquito cells. Virology 518:406–413.

Scolari F, Benoit JB, Michalkova V, Aksoy E, Takac P, Abd-Alla AMM, Malacrida AR, Aksoy S, Attardo GM. 2016. The Spermatophore in Glossina morsitans morsitans: Insights into Male Contributions to Reproduction. Sci Rep 6:20334.

Sharpe SR, Morrow JL, Brettell LE, Shearman DC, Gilchrist S, Cook JM, Riegler M. 2021. Tephritid fruit flies have a large diversity of co-occurring RNA viruses. Journal of Invertebrate Pathology:107569.

Sievers F, Higgins DG. 2014. Clustal Omega. Current Protocols in Bioinformatics 48:3.13.1-3.13.16.

Snyder AK, Deberry JW, Runyen-Janecky L, Rio RVM. 2010. Nutrient provisioning facilitates homeostasis between tsetse fly (Diptera: Glossinidae) symbionts. Proc Biol Sci 277:2389–2397.

Snyder AK, McLain C, Rio RVM. 2012. The Tsetse Fly Obligate Mutualist Wigglesworthia morsitans Alters Gene Expression and Population Density via Exogenous Nutrient Provisioning. Applied and Environmental Microbiology 78:7792–7797.

Snyder AK, Rio RVM. 2015. “Wigglesworthia morsitans” Folate (Vitamin B9) Biosynthesis Contributes to Tsetse Host Fitness. Applied and Environmental Microbiology 81:5375–5386.

Symula RE, Alam U, Brelsfoard C, Wu Y, Echodu R, Okedi LM, Aksoy S, Caccone A. 2013. Wolbachia association with the tsetse fly, Glossina fuscipes fuscipes, reveals high levels of genetic diversity and complex evolutionary dynamics. BMC Evolutionary Biology 13:31.

Telleria EL, Benoit JB, Zhao X, Savage AF, Regmi S, Silva TLA e, O’Neill M, Aksoy S. 2014. Insights into the Trypanosome-Host Interactions Revealed through Transcriptomic Analysis of Parasitized Tsetse Fly Salivary Glands. PLOS Neglected Tropical Diseases 8:e2649.

Thorvaldsdóttir H, Robinson JT, Mesirov JP. 2013. Integrative Genomics Viewer (IGV): high-performance genomics data visualization and exploration. Briefings in Bioinformatics 14:178–192.

Trappeniers K, Matetovici I, Van Den Abbeele J, De Vooght L. 2019. The Tsetse Fly Displays an Attenuated Immune Response to Its Secondary Symbiont, Sodalis glossinidius. Frontiers in Microbiology 10:1650.

Tsagmo Ngoune JM, Njiokou F, Loriod B, Kame-Ngasse G, Fernandez-Nunez N, Rioualen C, van Helden J, Geiger A. 2017. Transcriptional Profiling of Midguts Prepared from Trypanosoma/T. congolense-Positive Glossina palpalis palpalis Collected from Two Distinct Cameroonian Foci: Coordinated Signatures of the Midguts’ Remodeling As T. congolense-Supportive Niches. Front Immunol 8:876.

Tsagmo Ngoune JM, Reveillaud J, Sempere G, Njiokou F, Melachio TT, Abate L, Tchioffo MT, Geiger A. 2019. The composition and abundance of bacterial communities residing in the gut of Glossina palpalis palpalis captured in two sites of southern Cameroon. Parasites & Vectors 12:151.

Vigneron A, Aksoy E, Weiss BL, Bing X, Zhao X, Awuoche EO, O’Neill MB, Wu Y, Attardo GM, Aksoy S. 2018. A fine-tuned vector-parasite dialogue in tsetse’s cardia determines peritrophic matrix integrity and trypanosome transmission success. PLOS Pathogens 14:e1006972.

Wang J, Weiss B, Aksoy S. 2013. Tsetse fly microbiota: form and function. Frontiers in Cellular and Infection Microbiology 3:69.

Wang J, Wu Y, Yang G, Aksoy S. 2009. Interactions between mutualist Wigglesworthia and tsetse peptidoglycan recognition protein (PGRP-LB) influence trypanosome transmission. PNAS 106:12133–12138.

Weiss BL, Maltz M, Aksoy S. 2012. Obligate Symbionts Activate Immune System Development in the Tsetse Fly. The Journal of Immunology 188:3395–3403.

Weiss BL, Savage AF, Griffith BC, Wu Y, Aksoy S. 2014. The Peritrophic Matrix Mediates Differential Infection Outcomes in the Tsetse Fly Gut following Challenge with Commensal, Pathogenic, and Parasitic Microbes. The Journal of Immunology 193:773–782.

Weiss BL, Wang J, Aksoy S. 2011. Tsetse Immune System Maturation Requires the Presence of Obligate Symbionts in Larvae. PLOS Biology 9:e1000619.

Welburn SC, Maudlin I. 1992. The nature of the teneral state in Glossina and its role in the acquisition of trypanosome infection in tsetse. Ann Trop Med Parasitol 86:529–536.

Yang L, Weiss BL, Williams AE, Aksoy E, Orfano A de S, Son JH, Wu Y, Vigneron A, Karakus M, Aksoy S. 2021. Paratransgenic manipulation of a tsetse microRNA alters the physiological homeostasis of the fly’s midgut environment. PLOS Pathogens 17:e1009475.

Zulkower V, Rosser S. 2020. DNA Features Viewer: a sequence annotation formatting and plotting library for Python. Bioinformatics 36:4350–4352.

